## Supplemental Materials for "A Theory of Hippocampal Theta Correlations: Extrinsic and Intrinsic Sequences"

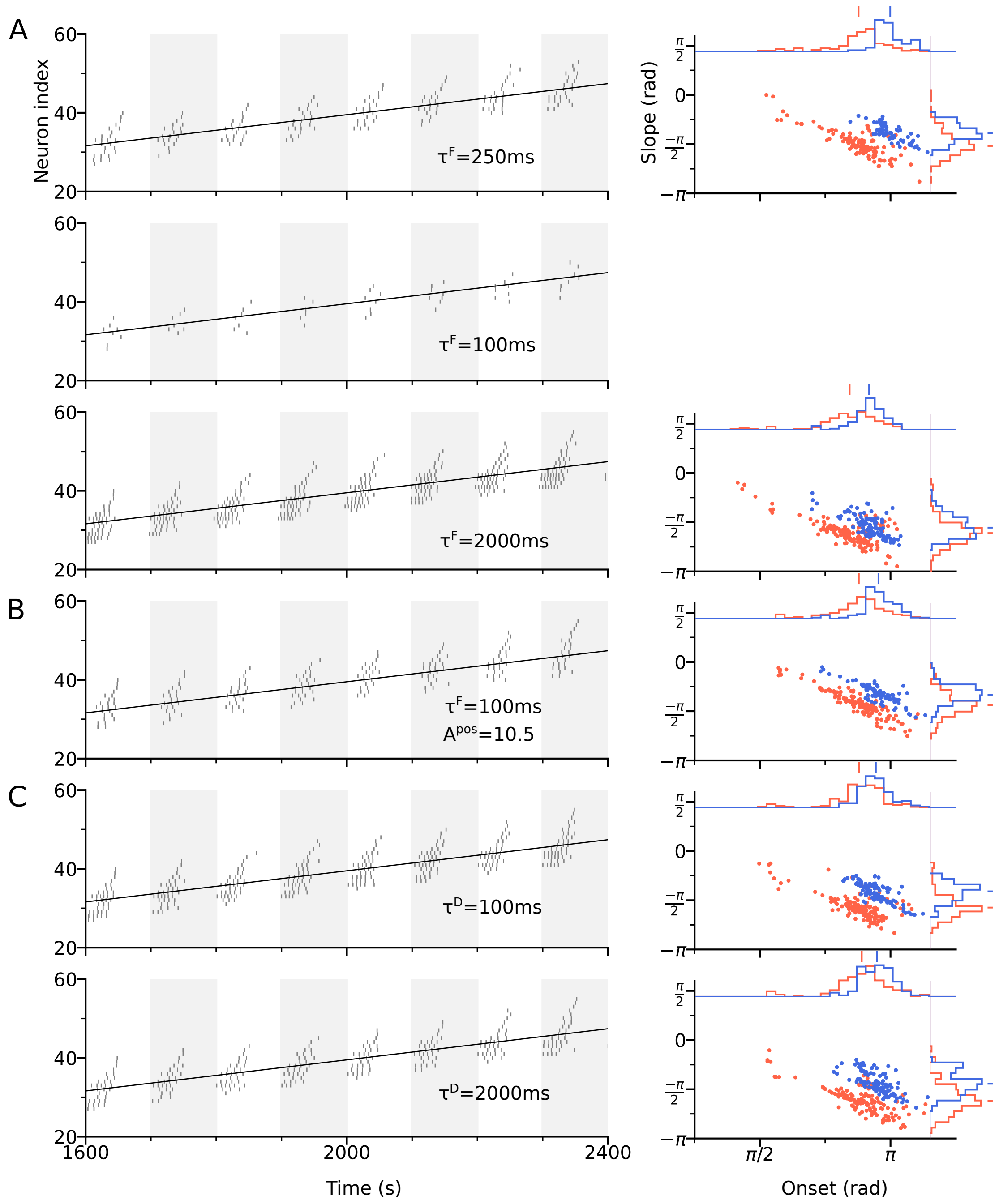


Figure 3—figure supplement 1: Effects of STF and STD time constants on theta sequences. (A) Left: Spike raster plot of the CA3 place cells along the trajectory at $\tau^F$=250ms (top row), 100ms (middle) and 2000ms (bottom). Conventions follow Figure 3B. Decreasing $\tau^F$ allows faster recovery time of the sensory input to its resting value $S_0^F = 0$, thus reducing the amount of depolarization and firing rate, as compared to $\tau^F$=500ms used in Figure 3. However, the temporal order of theta sequences remains unchanged. Right: Distribution of slopes and onsets of phase precession in the best and worst directions, following the conventions of Figure 3D. At $\tau^F$=100ms, the place cells do not generate enough spikes (spike count $\leq$ 5) for the analysis. (B) Increasing the amount of sensory input $A^{pos}$ by 60% (from 6.5 in Figure 3 to 10.5 here) can restore the firing activity and theta sequences at $\tau^F$=100ms. (C) Changing the STD time constant $\tau^D$ does not noticeably affect the temporal structure of theta sequences and the distribution of phase precession slopes and onsets.

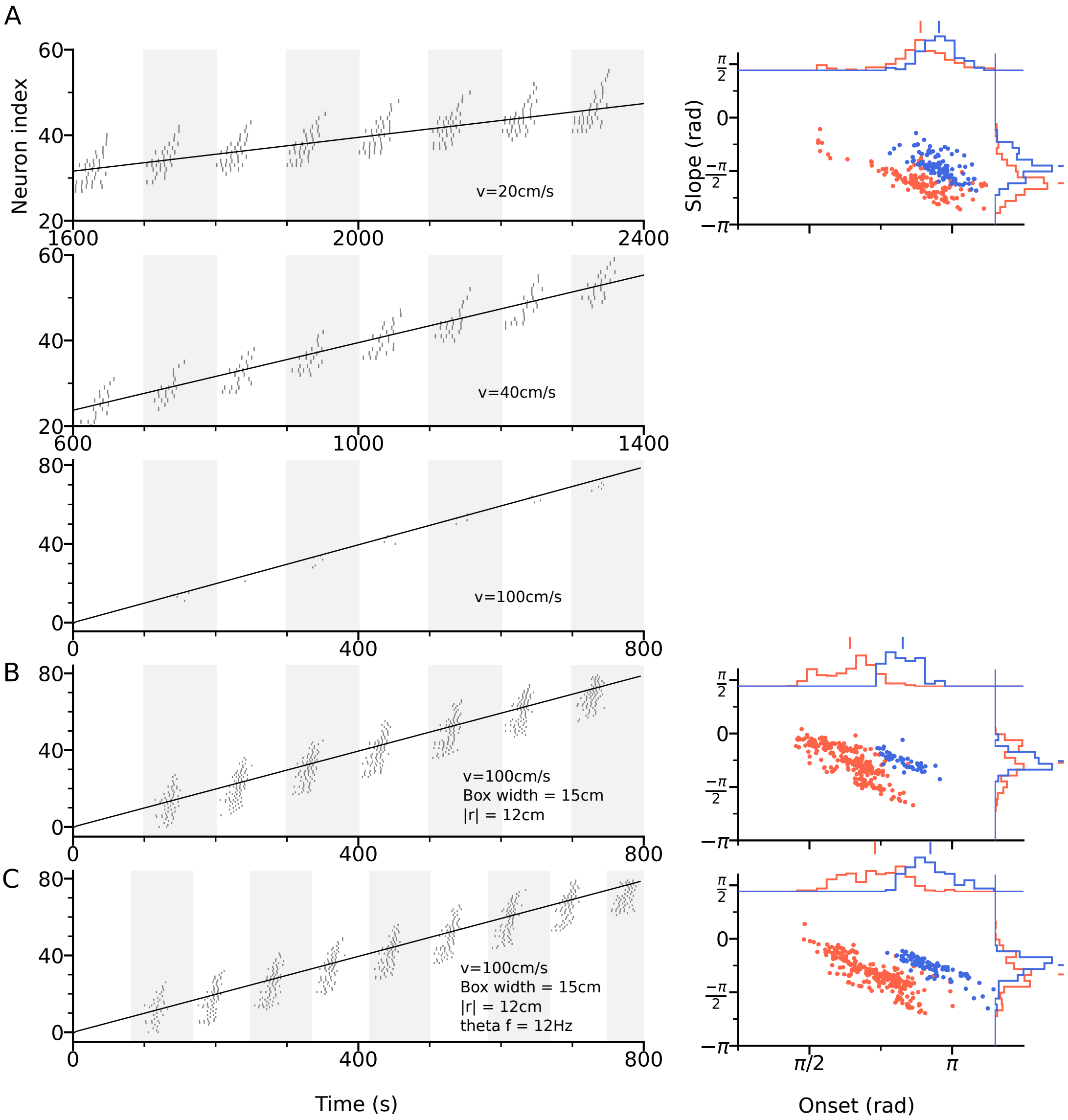


Figure 3—figure supplement 2: Effects of running speed on theta sequences. (A) Left: Spike raster plot of the CA3 place cells along the trajectory at running speed v=20cm/s (top row), 40cm/s (middle) and 100cm/s (bottom). Conventions follow Figure 3B. As place cells are traversed at increasing velocities, their firing rate decreases due to insufficient depolarization. Right: Distribution of slopes and onsets of phase precession in the best and worst directions, following the conventions of Figure 3D. Only the distribution for running speed at 20cm/s is shown. At running speed 40cm/s and 100cm/s, the place cells do not generate enough spikes (spike count $\leq$ 5) for the analysis. (B) Theta sequences at high running speed (100cm/s) could be produced by a larger place field size (width of box-shaped input is increased to 15cm compared to 5cm in Figure 3) and longer DG projection ($|\mathbf{r}|$=12cm compared to 4cm in Figure 3). Phase precession slopes and onsets are lower than in (A). (C) Increasing theta frequency to 12Hz can recover the decrease of phase precession slope and onset from the high running speed (Rivas et al., 1996; Maurer et al., 2005). Model parameters are the same as Figure 3 in the main text unless specified at the bottom right of the raster plots.

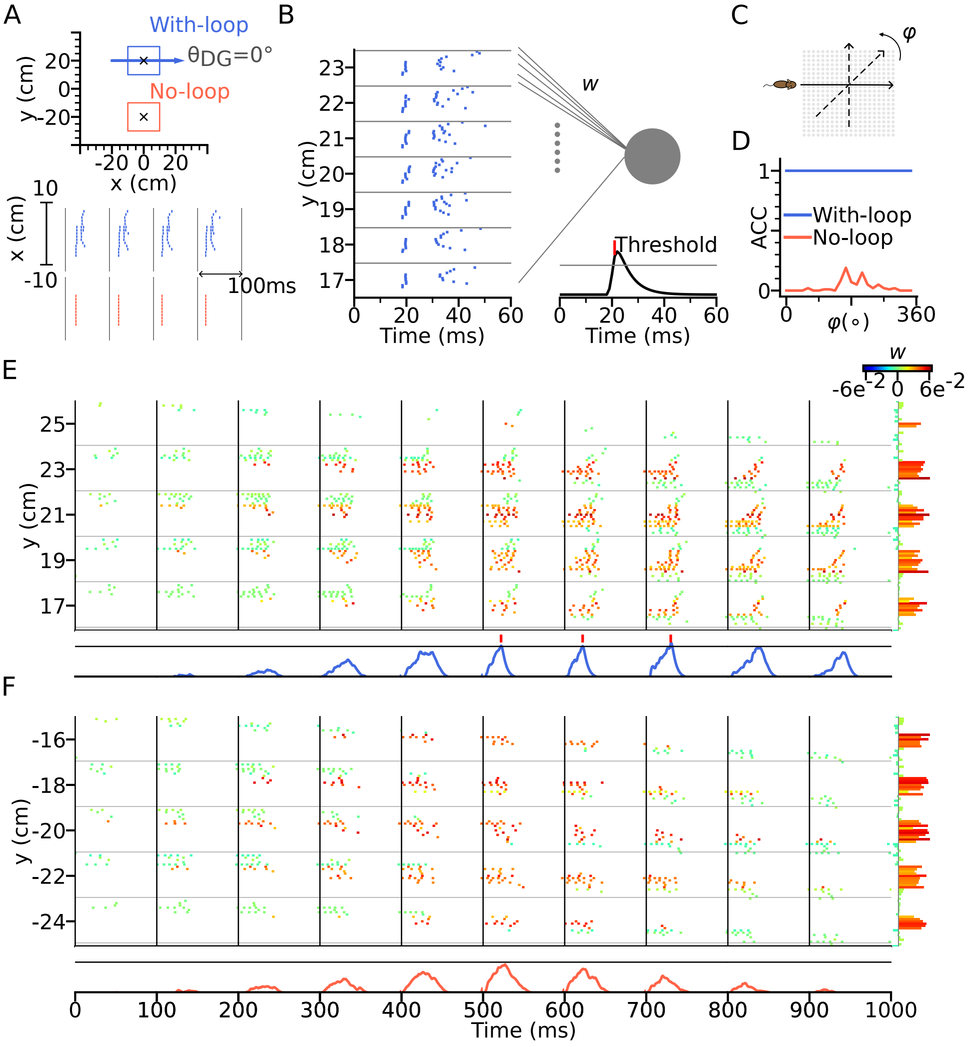


Figure 7—figure supplement 1: Decoding of positional landmarks using tempotrons in a network model without CA3-CA3 recurrence. Figure labels are the same as in Figure 7. Compared to the parameters in Figure 7, here CA3-CA3 weights were disabled ($B^{\mathrm{dir}} = 0$), but spatial input was increased during testing ($A^{\mathrm{dir}} = 14$, $S_0^F = 0.25$, $S_1^F = 1.5$) and DG-loop strengths were increased ($B^{\mathrm{DG}} = 5500$).
